# Pyrimethamine inhibits mutant NRF2 as a molecular glue

**DOI:** 10.1101/2025.08.18.670902

**Authors:** Zhaohui Xiong, Chorlada Paiboonrungruang, Haining Wang, Boopathi Subramaniyan, Candice Bui-Linh, Yahui Li, Huan Li, Francis Spitz, Xiaoxin Chen

## Abstract

Esophageal squamous cell carcinoma (ESCC) is a deadly disease and one of the most aggressive cancers of the gastrointestinal tract. As a master transcription factor regulating the stress response, nuclear factor erythroid 2-related factor 2 (NRF2) is often mutated, loses negative regulation by Kelch-like ECH-associated protein 1 (KEAP1), becomes hyperactive, and thus causes chemo-radioresistance and poor survival in human ESCC. Our previous research identified pyrimethamine (PYR) as an NRF2 inhibitor and demonstrated that inhibition of dihydrofolate reductase (DHFR) is the major mechanism of its action. Based on these findings, PYR has advanced into a Phase I window-of-opportunity clinical trial. In this study, using isogenic ESCC cell lines, we aimed to investigate the phenotypic consequences of NRF2 activation in ESCC cells and further elucidate the mechanism of action of PYR. While overexpression of wild-type NRF2 (NRF2^WT^) promoted squamous differentiation and suppressed proliferation, expression of mutant NRF2 (NRF2^Mut^) produced mixed effects on differentiation markers and promoted proliferation. NRF2 activation reduced sensitivity to chemotherapy and radiation, whereas PYR treatment restored chemosensitivity and radiosensitivity in NRF2^Mut^ ESCC cells. Co-immunoprecipitation and proximity ligation assays revealed that PYR selectively enhanced the interaction between NRF2^W24C^ and KEAP1, but not between NRF2^WT^ and KEAP1. Isothermal titration calorimetry (ITC) confirmed the direct binding of PYR to KEAP1 and surface plasmon resonance (SPR) showed that PYR modestly facilitated the interaction between a DLG^W24C^ peptide and the Kelch fragment. Molecular docking suggested that PYR bound to a pocket within the Kelch domain near Arg^415^. In summary, NRF2 activation promotes cell proliferation and therapy resistance in ESCC in a context-dependent manner. PYR functions as a modest molecular glue that selectively restores KEAP1 binding to NRF2^W24C^, providing a potential therapeutic strategy for NRF2^Mut^ ESCC.

## Introduction

Esophageal squamous cell carcinoma (ESCC) is a highly aggressive malignancy and ranks among the most lethal cancers of the gastrointestinal tract worldwide. Despite advances in surgery, radiation, and chemotherapy, the five-year survival rate for patients with advanced-stage ESCC remains dismal [1, 2]. One of the factors contributing to therapeutic resistance and poor prognosis in ESCC is hyperactivation of nuclear factor erythroid 2-related factor 2 (NRF2), a master transcription factor that regulates cellular defense mechanisms against oxidative stress, xenobiotics, and metabolic injury [3, 4]. Under physiological conditions, NRF2 activity is tightly controlled by its negative regulator, Kelch-like ECH-associated protein 1 (KEAP1), which targets NRF2 for ubiquitination and subsequent proteasomal degradation. This regulation is mediated through direct binding of two conserved motifs within NRF2 - the high-affinity ETGE motif and the lower-affinity DLG motif - to the Kelch domain of KEAP1, enabling proper positioning of NRF2 for ubiquitination by the CUL3-based E3 ligase complex [5]. However, in approximately 10-22% of human ESCC cases, this regulatory axis is disrupted by gain-of-function mutations in NRF2 (NRF2^Mut^) or loss-of-function mutations in KEAP1, leading to persistent NRF2 stabilization and nuclear accumulation [6–8]. Although NRF2 plays a protective role in normal tissues by mitigating oxidative stress, its sustained activation in cancer cells confers a survival advantage, enhances proliferation, rewires metabolic pathways, and drives resistance to chemotherapy, radiation, and immunotherapy [5]. These observations underscore NRF2 as a compelling therapeutic target, especially in cancers like ESCC where mutations affecting the NRF2-KEAP1 interaction are common.

Despite its importance, targeting NRF2 pharmacologically has proven challenging, as it is considered an ’undruggable’ target due to its function as a transcription factor lacking classical enzymatic pockets or well-defined ligand-binding sites [9, 10]. Nonetheless, significant progress has been made in recent years. A notable example is VVD-065, a molecular glue that specifically and covalently engages Cys^151^ on the BTB domain of KEAP1, which in turn promotes KEAP1-CUL3 interaction, leading to enhancement of degradation of both wild-type NRF2 (NRF2^WT^) and NRF2^Mut^. While VVD-065 has shown promise, its activity is limited to certain NRF2 mutations [11]. A related compound, VVD-130037, which employs the same molecular glue mechanism, is currently undergoing Phase 1 clinical trial (NCT05954312).

Another promising agent is pyrimethamine (PYR), an FDA-approved drug long used to treat malaria and toxoplasmosis by inhibiting dihydrofolate reductase (DHFR) [12]. We identified PYR as a novel NRF2 inhibitor through small-molecule screening and validated its efficacy both *in vitro* and *in vivo* [13]. PYR was found to suppress NRF2 expression and attenuate *Nrf2^E79Q^*-driven esophageal hyperplasia in a mouse model. Importantly, it achieved these effects at clinically relevant and safe doses, making it an ideal candidate for drug repurpose. In support of its translational potential, PYR has progressed to a Phase 1 clinical trial (NCT05678348). Genetic, pharmacologic, and metabolic epistasis studies demonstrated that DHFR inhibition was essential for the NRF2-inhibitory activity of PYR and its structural analogue WCDD115. Proteomic analyses showed that PYR shares a molecular signature with methotrexate (MTX), a classical DHFR inhibitor [14]. On the other hand, in NRF2^W24C^-KYSE70 and NRF2^D77V^-KYSE180 cells, PYR promoted NRF2^Mut^ ubiquitination and proteasomal degradation and shortened its half-life, suggesting an additional mode of action as a molecular glue degrader [13]. Interestingly, similar protein degradation effect of PYR was observed with the exon 2-depleted splice variant of aminoacyl-tRNA synthetase-interacting multifunctional protein 2 (AIMP2-DX2) in lung cancer [15]. These findings underscore the potential of PYR as a dual-function NRF2 inhibitor, acting through DHFR inhibition and direct NRF2 degradation. The current study aims to further dissect PYR’s activity as a molecular glue, which may inform the development of more potent and specific inhibitors for NRF2^Mut^ malignancies including ESCC.

### Materials and Methods Cell culture and chemicals

Human ESCC cells, KYSE70 (NRF2^W24C^), KYSE180 (NRF2^D77V^), and KYSE450 (NRF2^WT^) were purchased from DSMZ (Braunschweig, Germany). TE14 (NRF2^D29H^) and OE21 (NRF2^G81S^) cells were obtained from Columbia University (New York, NY) and Sigma-Aldrich (St. Louis, MO), respectively. All cells were cultured in Gibco RPMI1640 (ThermoFisher, Waltham, MA) supplemented with 10% FBS and 1% antibiotics (penicillin/streptomycin) at 37°C under humidified conditions with 5% CO2. PYR, MTX, 5-fluorouracil (5-FU) and cisplatin were purchased from Sigma-Aldrich. NRF2^null^-KYSE70, KEAP1^null^-KYSE70, and KEAP1^null^-KYSE450 cells were generated by using CRISPR-Cas9-mediated genome editing.

### RT-PCR

Total RNA was isolated using Qiagen RNeasy Mini Kit (Germantown, MD) and reverse transcribed using the Advantage RT-for-PCR kit (Clontech, Mountain View, CA), according to the manufacturer’s instructions. mRNA transcripts were detected with the following primer pair: forward (5’-TCATGATGGACTTGGAGCTG-3’) and reverse (5’-GCAATGAAGACTGGGCTCTC-3’). This primer pair was expected to amplify all mRNA transcripts including transcript 1 (475bp), transcript 6 (385bp), and transcript 7 (256bp) [16].

### Western blotting and co-immunoprecipitation (co-IP)

The whole cell lysates were made using RIPA lysis buffer supplemented with a protease inhibitor cocktail (ThermoFisher). Proteins were separated by SDS-PAGE and transferred to nitrocellulose membranes. Membranes were blocked and then incubated with a primary antibody overnight at 4°C (Table S1). Chemiluminescence was detected using iBright FL1500 (ThermoFisher) and followed by quantification using ImageJ. To analyze NRF2-KEAP1 binding, cells were treated with PYR or MTX for 2 h before being harvested in RIPA buffer. The cell lysate was pre-cleaned with protein G agarose beads (Roche Diagnostic, Basel, Switzerland) for 1 h at 4°C. Samples were incubated with 1 µg anti-NRF2 or anti-KEAP1 antibody overnight at 4°C with rotation, and then 25 µl protein G agarose beads was added for 1.5 h at 4°C. Immunoprecipitants were washed and boiled in 2X SDS loading buffer at 95 °C for 5 min, and analyzed by Western blotting.

### Cell cycle analysis

Cells were harvested by trypsinization, washed with PBS, and fixed in 70% ethanol. Fixed cells were stored at -20°C overnight. Cells were washed and resuspended in PBS containing RNase A, filtered, and incubated with propidium iodide (50 µg/ml) for 1 h at 4°C in the dark. Cell cycle analysis was performed using the BD FACSymphony™ A3 Cell Analyzer and FlowJo™ v10 software (BD Biosciences, Milpitas, CA).

### Proliferation assay

Cells were seeded in 96-well plates and incubated with NucRed™ Live 647 ReadyProbes™ Reagent (ThermoFisher) to label nuclei. Plates were imaged every 4 h over a 72-h period using the Incucyte® SX5 Live-Cell Analysis System (Sartorius, Göttingen, Germany). Fluorescent images were analyzed using the Incucyte software, and proliferation curves were generated using GraphPad Prism.

### Apoptosis assay

Apoptosis was assessed using the CellEvent™ Caspase-3/7 Green Detection Reagent (ThermoFisher). Cells were seeded in 96-well plates and treated as indicated. Caspase-3/7 reagent was added according to the manufacturer’s instructions. Plates were imaged every 4 h for 72 h using the ImageXpress Pico Automated Cell Imaging System (Molecular Devices, San Jose, CA). Green fluorescence indicating caspase-3/7 activation was quantified using the system’s integrated analysis software.

### Chemosensitivity assay

To evaluate the drug effects on cell viability, real-time imaging was performed using the IncuCyte platform. KYSE70 and NRF2^null^-KYSE70 cells were seeded in 96-well plates at a density of 1,000 cells per well and incubated overnight to allow for cell adherence. Cells were pretreated with 10 µM PYR for 48 h. Subsequently, either 5-FU or cisplatin was administered to assess differential chemosensitivity. 5-FU was serially diluted in a 1:2 ratio from an initial concentration of 1,000 µM, yielding a total of 11 concentrations plus a vehicle control (0 µM). Cisplatin was prepared using the same dilution scheme starting from 240 µM. To facilitate real-time monitoring of cell viability, NucRed™ Live 647 ReadyProbes™ Reagent (ThermoFisher) was utilized to stain nuclei, while YOYO-1 iodide (ThermoFisher) was employed as a cell-impermeant dye to selectively identify dead cells. Following treatment and dye addition, plates were transferred to the Incucyte platform. Phase-contrast and fluorescence images were acquired at 6-h intervals over a period of 48 h. Cell confluence and fluorescence signals were quantified using the Incucyte integrated analysis software.

### Radiosensitivity assays

Cells were seeded into a 3D pillar plate format on Day 0 using the MBD ASFA® Spotter (Medical & Bio Decision, Korea), which enables precise, non-contact dispensing of cell-laden hydrogel droplets. Each spot contained 1,000 cells embedded in a 3D matrix to support spheroid formation. On Day 1 and Day 4, the cultured cells were exposed to varying doses of ionizing radiation to assess radiosensitivity. On Day 7, cell viability was evaluated using the CellTiter-Glo^®^ 3D Cell Viability Assay (Promega, Madison, WI) following the manufacturer’s instructions.

The effect of PYR in combination with ionizing radiation on cell viability was also assessed in 2D-cultured cells. Cells were seeded into 96-well plates at a density of 2,500 cells per well and incubated overnight to allow for adherence. Cells were pretreated with 10 µM PYR for 48 h, followed by exposure to graded doses of X-ray irradiation (0, 2, 4, 6, 8, and 10 Gy) using an irradiator (RS2000; RadSource, Brentwood, TN). After irradiation, cells were incubated for an additional 72 h under standard culture conditions. Cell viability was assessed using the CellTiter-Glo^®^ Luminescent Cell Viability Assay following the manufacturer’s instructions. Luminescence was measured using a microplate reader SpectraMax® iD5 (Molecular Devices, San Jose, CA), and viability was normalized to the non-irradiated control.

### Metabolomic profiling

KYSE70 and NRF2^null^-KYSE70 cell pellets were analyzed by Metabolon (Durham, NC) with ultra-high performance liquid chromatography-tandem mass spectroscopy. Raw data were extracted, peak-identified and QC processed by Metabolon.

### RNAseq

RNAseq was performed by Admera Health (South Plainfield, NJ). Total RNA was used as input material for the RNA sample preparations. Sequencing libraries were generated using NEBNext Ultra TM RNA Library Prep Kit for Illumina (NEB, Ipswich, MA) following the manufacturer’s recommendations and index codes were added to attribute sequences to each sample. After library preparation, the samples were sequenced (150 bp) according to manufacturer specifications. The quality-filtered reads were aligned with STAR (version 2.6.90c) to the human reference genome (hg38) with its respective RefSeq annotation, and the expression levels of genes were obtained with Feature Counts (version 1.5.1). DESeq2 (version 1.34.1) was used to identify the differentially expressed genes (DEGs). Data matrices were normalized and the DEGs were reported according to the fold change cut-off and corrected modeling p-values. Gene set enrichment analysis was conducted to evaluate the differential enrichment of gene sets including “NRF2 target gene set”. The raw data has been deposited in NCBI GEO database (GSE 299159 and GSE 299160).

### Proximity ligation assay (PLA)

The Duolink In Situ Red Fluorescent kit (Sigma-Aldrich) was employed to investigate the endogenous interaction between NRF2 and KEAP1 in human ESCC cells. The PLA was conducted following the manufacturer’s instructions. PLA with anti-NRF2 and anti-KEAP1 alone served as negative controls. A total of 1 x 10^6^ cells were plated on poly L-lysine coated coverslips and allowed to adhere for 24 h. The cells were treated with PYR for 4 h and fixed using 4% paraformaldehyde and permeabilized with 0.1% Triton X-100 before being incubated with the antibodies. The nuclei were counterstained with DRAQ5 (ThermoFisher). Images were captured with a confocal microscope and processed with Leica Application Suite X. PLA signals in a minimum of 100 cells were counted using ImageJ.

### Isothermal titration calorimetry (ITC)

ITC assay was performed by Creative Biolabs (Shirley, NY). 4 μl aliquots of 100 μM PYR were injected from a syringe into the sample cell containing 200 μl of human recombinant KEAP1 (MedChemExpress, Monmouth Junction, NJ). Each experiment was accompanied by the corresponding control experiment in which 4 μl aliquots of 100 μM PYR were injected into a buffer alone. The delay between injections was 150 s. Each injection generated a heat burst curve. Calorimetric titrations were performed at 25 °C with a speed of 250 rpm.

### Surface plasmon resonance (SPR)

A Kelch fragment (Ala^321^-Glu^611^ of human KEAP1 protein, MW 34,038.03 Da) was expressed *in vitro*, purified, biotinylated, and validated by LC-MS. DLG^WT^ (^17^MDLIDILWRQDIDLGVSREVFDFS^40^) and DLG^W24C^ (^17^MDLIDILCRQDIDLGVSREVFDFS^40^) were synthesized. SPR analysis was performed using Biacore S200 (Cytiva, Marlborough, MA) equipped with a streptavidin sensor chip. Biotinylated Kelch (4.05 mg/ml) was immobilized on flow cells, respectively, at a concentration of 10 μg/ml, with a flow rate of 5 μl/min for 44 or 49 s. Analytes (DLG^WT^ and DLG^W24C^) were prepared as 2-fold serial dilutions ranging from 200 μM to 0.39 μM, while PYR was diluted from 500 μM to 0.97 μM. Each analyte was injected at a flow rate of 30 μl/min with an association phase of 90 s and a dissociation phase of 180 s. The sample compartment was maintained at 25 °C, and the analysis was conducted at 15 °C.

### Molecular docking

DiffDock was used to explore potential PYR-binding sites on KEAP1. We further generated NRF2 structure with AlphaFold 2 (DeepMind, London, UK) and then docked PYR on KEAP1 (PDB: 4IFJ) and NRF2 using MOE (Chemical Computing Group, Montreal, Canada) employing rigid docking with the Amber10 force field to generate the docking models. Interaction between KEAP1 and three NRF2 activators which disrupts NRF2-KEAP1 interaction, PRL-295 [17], KI696 [18], and Compound 4 [19], were visualized to demonstrate the differences between PYR and these activators.

### Statistical analysis

GraphPad Prism 10 (GraphPad Software, La Jolla, CA) was used two-way ANOVA and Student’s t-test, with the statistical significance level set at 0.05. Drug response data were processed to generate dose-response curves and to calculate IC□□ using GraphPad Prism. Statistical comparisons of drug response were performed using two-way ANOVA. For metabolomic profiling, refined data were analyzed by two-way ANOVA and plotted with GraphPad Prism. To demonstrate the synergistic effects, combination index (CI) analysis was performed using the CompuSyn software (CompuSyn Inc, Paramus, NJ). This program defines synergism as an effect that exceeds the expected additive effect with a CI < 1.

## Results

### Establishment of isogenic ESCC cells with NRF2 or KEAP1 deficiency

To investigate the molecular and phenotypic consequences of NRF2 activation in ESCC, we generated isogenic human ESCC cell lines via CRISPR-Cas9 gene editing. Monoclonal NRF2^null^-KYSE70, KEAP1^null^-KYSE70, and KEAP1^null^-KYSE450 cells were established and compared with their respective parental lines, NRF2^W24C^-KYSE70 and NRF2^WT^-KYSE450. These isogenic cell lines were validated through Western blotting, Sanger sequencing, RNA-seq, and untargeted metabolomics (Figures S1, S2, and 1). As expected, RNAseq analysis demonstrated that the “human ESCC NRF2 target gene set” was significantly enriched in NRF2^W24C^-KYSE70 and KEAP1^null^-KYSE450 cells relative to their isogenic cells (Excel S1 and S2). In line with NRF2’s established role in metabolic regulation, metabolomic profiling revealed elevated levels of glutathione metabolism, glycolysis, gluconeogenesis, pyruvate metabolism, and phospholipid-related pathways in NRF2^W24C^-KYSE70 cells compared to NRF2^null^-KYSE70 cells (Excel S3). However, a faint and smaller NRF2 band was detected by Western blotting in NRF2^null^-KYSE70 cells (Figure 1A). Given that human NRF2 mRNA has eight transcripts [20], RT-PCR analysis showed expression of variant 6 (encoding protein isoform 4) and variant 7 (encoding protein isoform 5) in NRF2^null^-KYSE70 cells (Figure S1C). Since protein isoform 4 is ∼3kD smaller and protein isoform 5 is ∼8 kD smaller than the predominant protein isoform 1, the smaller NRF2 band observed on Western blot is believed to be protein isoform 5.

**Figure 1.**
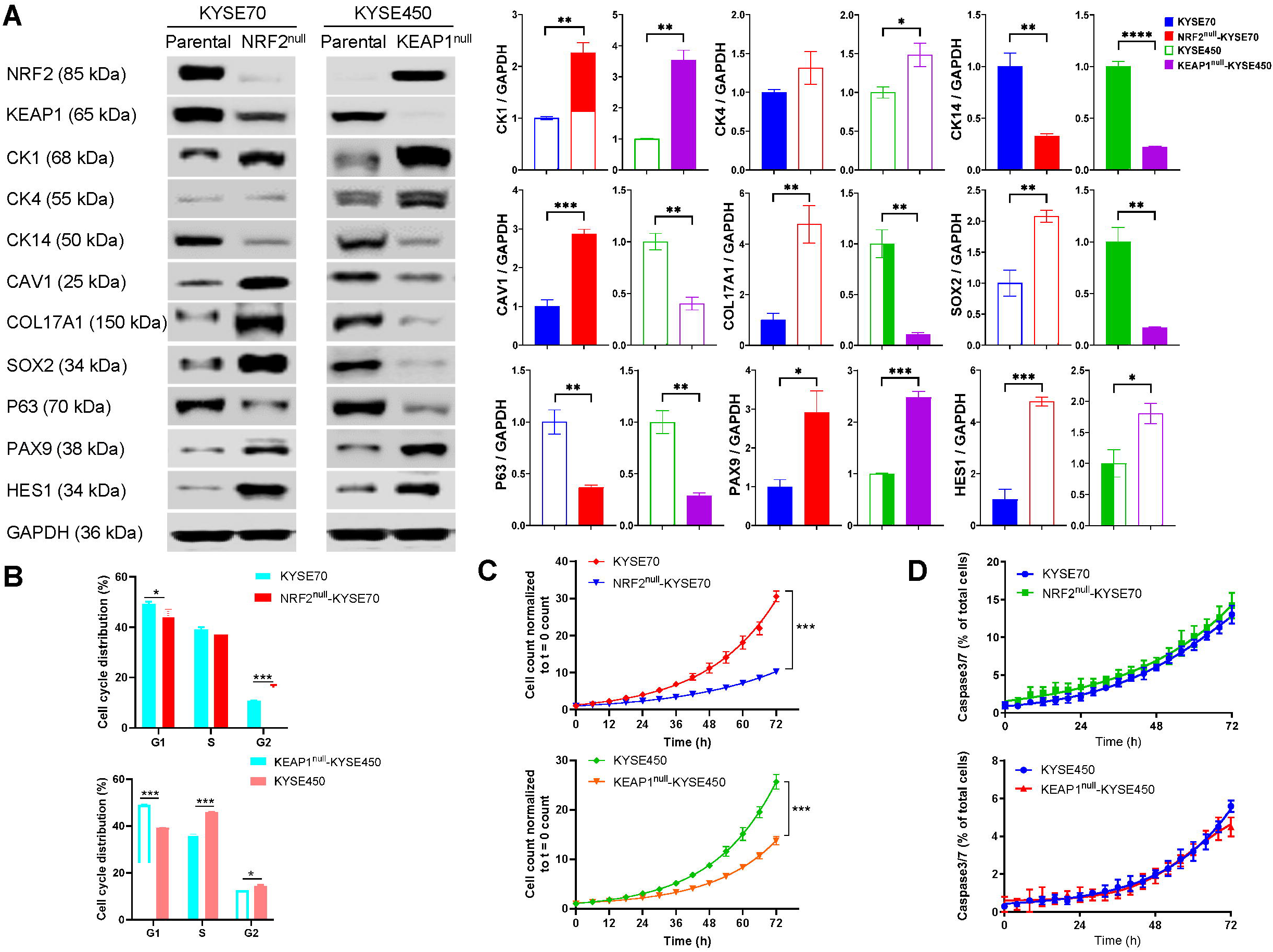
NRF2 modulated gene expression and cellular behaviors of human ESCC cells. NRF2^null^-KYSE70 cells were compared with NRF2^W24C^-KYSE70 cells, and KEAP1^null^-KYSE450 cells with NRF2^WT^- KYSE450 cells, on the following parameters: (A) Squamous differentiation markers as detected by Western blotting; (B) Cell cycle distribution as detected by cell sorting; (C) Cell proliferation; and (D) Cell apoptosis as detected by IncuCyte live cell imaging. *\* p* < 0.05, ** *p* < 0.01, *\*\*\* p* < 0.001, **** *p* < 0.0001. Bar represented mean ± SD.

### NRF2 activation modulates cellular behaviors, chemosensitivity and radiosensitivity of human ESCC cells

RNAseq analysis revealed that both the “basal layer gene set” and “P63 target gene set” were enriched in KYSE450 cells in comparison to KEAP1^null^-KYSE450 cells, indicating that NRF2^WT^ overexpression promoted squamous differentiation (Excel S1, S2). Consistent with the transcriptomic data, Western blotting demonstrated that NRF2^WT^ overexpression upregulated squamous differentiation markers (CK1, CK4) and NOTCH target genes (PAX9, HES1), while downregulating basal cell markers (CK14, CAV1, COL17A1, SOX2, P63) (Figure 1A). In contrast, NRF2^W24C^ expression exerted differential effects: it downregulated squamous differentiation markers (CK1) and NOTCH target genes (PAX9, HES1) but had a mixed effect on basal cell markers—downregulating CAV1, COL17A1, and SOX2, while upregulating CK14 and P63. These results suggest that NRF2^W24C^ may have distinct transcriptional outputs from NRF2^WT^, particularly in its regulation of squamous differentiation.

We further investigated the impact of NRF2 activation on cell cycle distribution, proliferation, and apoptosis. Flow cytometry analysis revealed that NRF2 activation altered cell cycle distribution in both KYSE70 and KYSE450 cells (Figure 1B). Live-cell imaging demonstrated that NRF2^W24C^ expression enhanced cell proliferation in KYSE70 cells, whereas NRF2^WT^ overexpression inhibited proliferation in KYSE450 cells (Figure 1C). However, neither form of NRF2 significantly affected apoptosis in either cell line (Figure 1D). These findings suggest that NRF2 activation modulates key cellular behaviors in human ESCC cells, with NRF2^WT^ and NRF2^Mut^ exhibiting distinct functional effects.

To explore the role of NRF2 activation in modulating chemosensitivity in human ESCC cells, we treated isogenic NRF2^null^-KYSE70 and NRF2^W24C^-KYSE70 cells with 5-FU or cisplatin for 48 h. As expected, 5-FU markedly reduced the viability of NRF2^null^-KYSE70 cells (IC□□ = 50.53 µM), in comparison to NRF2^W24C^- KYSE70 cells (IC□□ = 79.28 µM). A similar trend was observed with cisplatin, supporting the notion that NRF2^W24C^ expression promotes chemoresistance in KYSE70 cells (Figure 2A).

**Figure 2.**
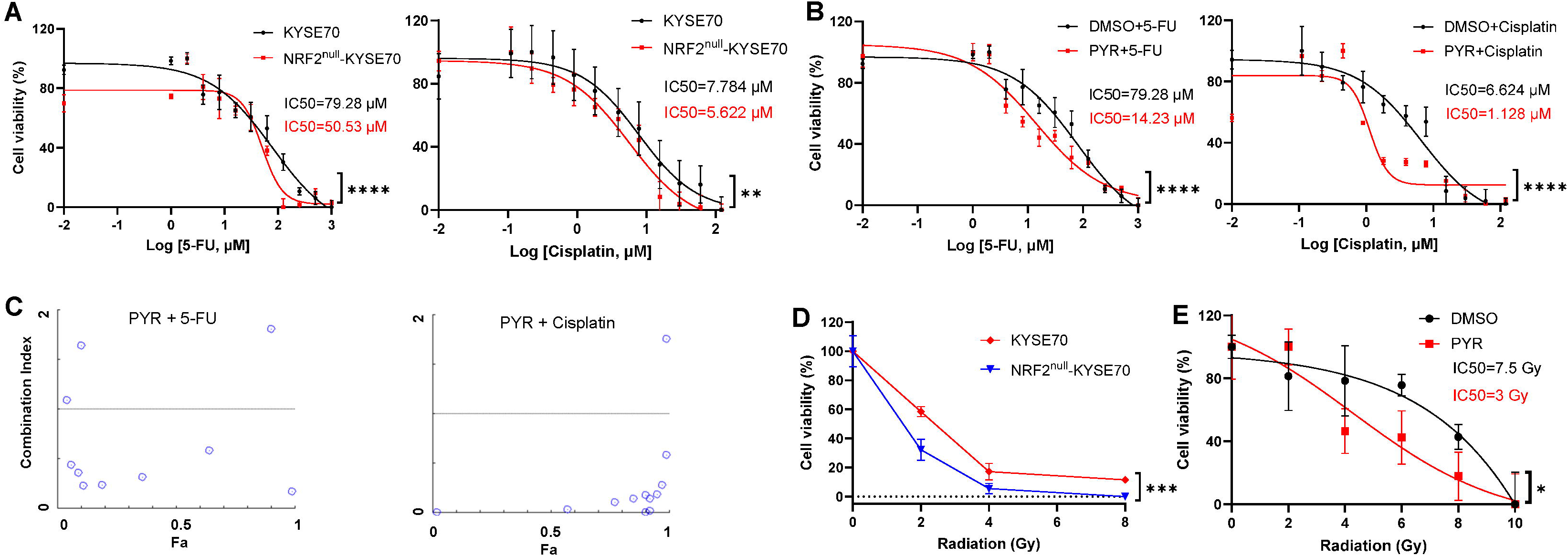
NRF2 modulated chemosensitivity and radiosensitivity of NRF2^W24C^-KYSE70 cells. (A) Viability of NRF2^W24C^-KYSE70 and NRF2^null^-KYSE70 cells after 5-FU or cisplatin treatment; (B) Viability of NRF2^W24C^- KYSE70 cells after co-treatment with PYR (10 μM) and 5-FU or cisplatin; (C) CI of PYR and 5-FU or cisplatin for NRF2^W24C^-KYSE70 cells; (D) Viability of NRF2^W24C^-KYSE70 and NRF2^null^-KYSE70 cells in 3D culture when exposed to radiation; (E) Viability of NRF2^W24C^-KYSE70 treated with PYR (10 μM) and radiation in 2D culture. *\* p* < 0.05, ** *p* < 0.01, *\*\*\* p* < 0.001, **** *p* < 0.0001. Bar represented mean ± SD.

In our previous work, we identified PYR as a small-molecule NRF2 inhibitor and validated its ability to suppress NRF2 expression both *in vitro* and *in vivo* in a NRF2^E79Q^-driven mouse model of esophageal hyperproliferation and hyperkeratinization [13]. In this study, co-treatment with PYR significantly enhanced the efficacy of 5-FU in NRF2^W24C^-KYSE70 cells, lowering the IC□□ from 79.28 µM to 14.23 µM. Similarly, co-treatment with PYR reduced the IC□□ of cisplatin from 6.624 µM to 1.128 µM (Figure 2B). Comparable chemosensitizing effects were also observed in NRF2^D77V^-KYSE180 cells treated with PYR in combination with either 5-FU or cisplatin (Figure S3A). Moreover, CI analysis demonstrated a synergistic interaction between PYR and both chemotherapeutic agents in KYSE70 cells, as indicated by CI values below 1 (Figure 2C). NRF2^null^-KYSE70 cells also exhibited increased radiosensitivity compared to the parental KYSE70 cells when cultured in 3D conditions (Figure 2D). In contrast, KEAP1^null^-KYSE450 cells showed enhanced radioresistance relative to parental KYSE450 cells (Figure S3B). Notably, co-treatment with PYR significantly sensitized NRF2^W24C^-KYSE70 cells to radiation in 2D culture, reducing the IC□□ from 7.5 Gy to 3 Gy (Figure 2E). Collectively, these findings indicate that NRF2 activation reduces both chemosensitivity and radiosensitivity in human ESCC cells, whereas pharmacological inhibition of NRF2 with PYR restores sensitivity to chemotherapy and radiation.

### PYR promotes NRF2^W24C^ degradation by enhancing NRF2^W24C^-KEAP1 binding

Our previous study has shown that PYR reduced NRF2^Mut^ half-life by promoting its ubiquitination and degradation in NRF2^W24C^-KYSE70 and NRF2^D77V^-KYSE180 cells, but not in NRF2^WT^-KYSE450 cells [13]. Similarly, PYR has been reported to promote ubiquitination and degradation of AIMP2-DX2 in lung cancer cells [15]. These data suggest that PYR may also act through the ubiquitin-proteasomal degradation pathway. In this study, PYR treatment for 4 h was found to inhibit NRF2^W24C^ expression in KYSE70 cells in a dose-dependent manner (Figure 3A). This inhibitory effect on NRF2^W24C^ was dependent on the presence of KEAP1, as PYR did not suppress NRF2^W24C^ expression in KEAP1^null^-KYSE70 cells (Figure 3B). Interestingly, this degradation effect on NRF2 in 4 h was dependent on specific point mutations, as PYR slightly reduced NRF2 expression in NRF2^D77V^-KYSE180 cells (Figure S3A), but had no effect in NRF2^WT^-KYSE450, NRF2^D29H^-TE14, or NRF2^G81S^-OE21 cells (Figure S3B, C, D).

**Figure 3.**
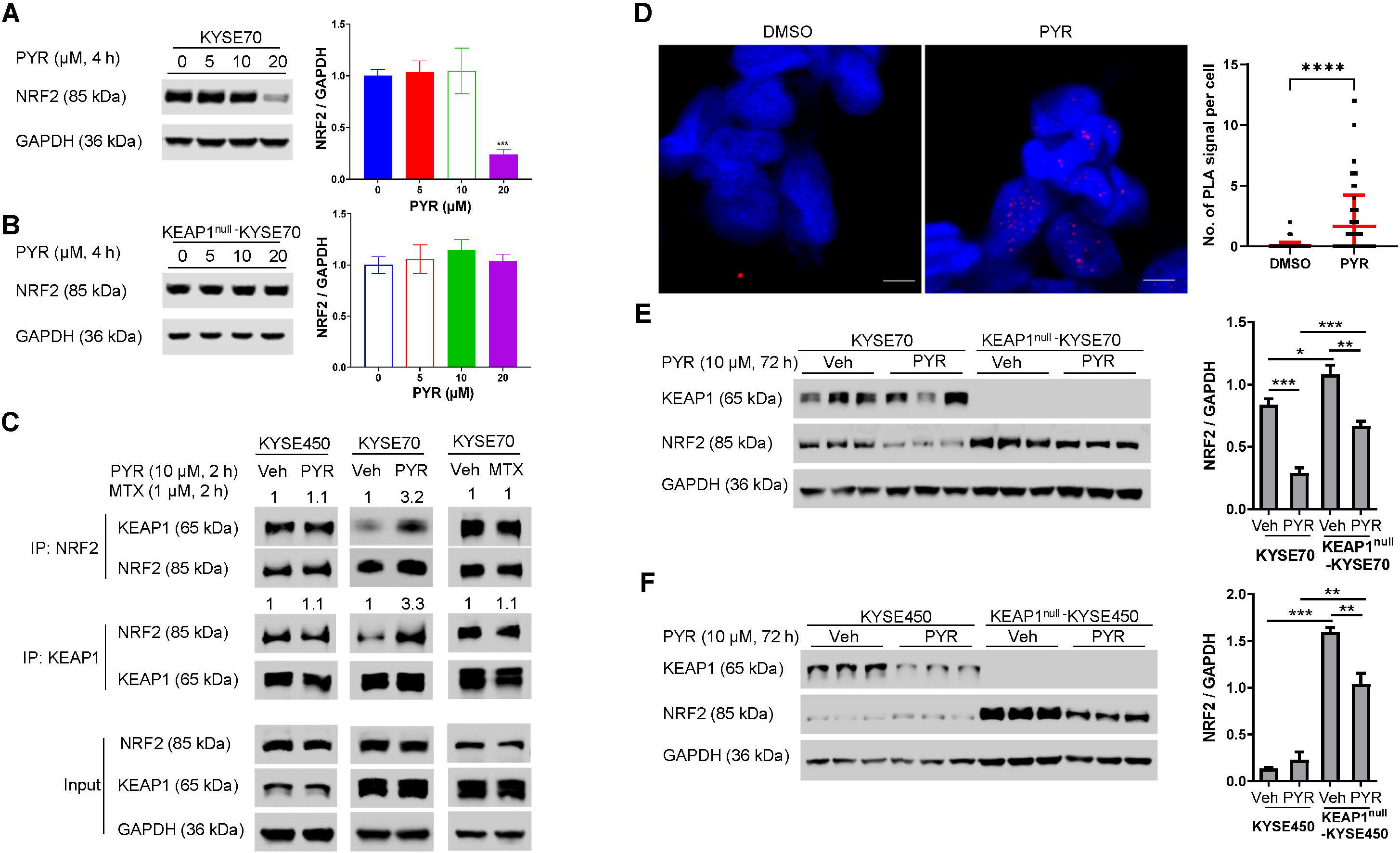
PYR promoted NRF2^W24C^-KEAP1 binding and NRF2^W24C^ degradation. (A) PYR degraded NRF2^W24C^ in KYSE70 cells in a dose-dependent manner in 4 h. (B) PYR failed to degrade NRF2^W24C^ in KEAP1^null^-KYSE70 cells in 4 h. (C) PYR (10 μM, 2 h) promoted NRF2^W24C^-KEAP1 interaction in KYSE70 cells, but not NRF2^WT^-KEAP1 interaction in KYSE450 cells, as shown by Co-IP. MTX (1 μM, 2 h) also failed to promote NRF2^W24C^-KEAP1 interaction in KYSE70 cells. (D) PYR (10 μM, 4h) increased the number of PLA signals in NRF2^W24C^-KYSE70 cells. (E) PYR (10 μM, 72 h) downregulated NRF2 expression in KEAP1^null^-KYSE70 cells as in NRF2^W24C^-KYSE70 cells; (F) PYR (10 μM, 72 h) downregulated NRF2 expression in KEAP1^null^-KYSE450 cells. *\* p* < 0.05, *\*\* p* < 0.01, *** *p* < 0.001, *\*\*\*\* p* < 0.0001. Bar represented mean ± SD.

Co-IP using anti-NRF2 or anti-KEAP1 revealed that PYR (10 μM, 2 h) enhanced the interaction between NRF2^W24C^ and KEAP1, but not between NRF2^WT^ and KEAP1 (Figure 3C). Treatment with another DHFR inhibitor, MTX, did not affect the NRF2^W24C^-KEAP1 interaction, indicating a specific effect of PYR. PLA analysis further confirmed that PYR significantly increased PLA signals in KYSE70 cells (Figure 3D), validating PYR’s role in enhancing NRF2^W24C^-KEAP1 interaction.

To further investigate whether the NRF2-inhibitory action of PYR depends on KEAP1 in a long term (72 h), we treated NRF2^W24C^-KYSE70 cells and NRF2^WT^-KYSE450 cells with or without KEAP1 with PYR (10 μM) and examined the expression of NRF2 in these cells. Contrary to the data of 4-h treatment (Figure 3A, B), 72-h treatment with PYR inhibited NRF2 expression in the absence of KEAP1, albeit to a lesser degree (Figure 3E, F).

To further validate the glue activity of PYR, SPR assay was established using two 24-mer peptides (DLG^WT^ and DLG^W24C^) and a Kelch fragment of KEAP1 (Ala^321^-Glu^611^). As expected, SPR analysis showed that DLG^WT^ was bound to the immobilized Kelch fragment (Figure 4A), while DLG^W24C^ exhibited no detectable binding (Figure 4B). PYR slightly increased the binding affinity between DLG^W24C^ and Kelch (Figure 4C, D), suggesting that PYR functions as a modest glue to stabilize the NRF2^W24C^-KEAP1 interaction.

**Figure 4.**
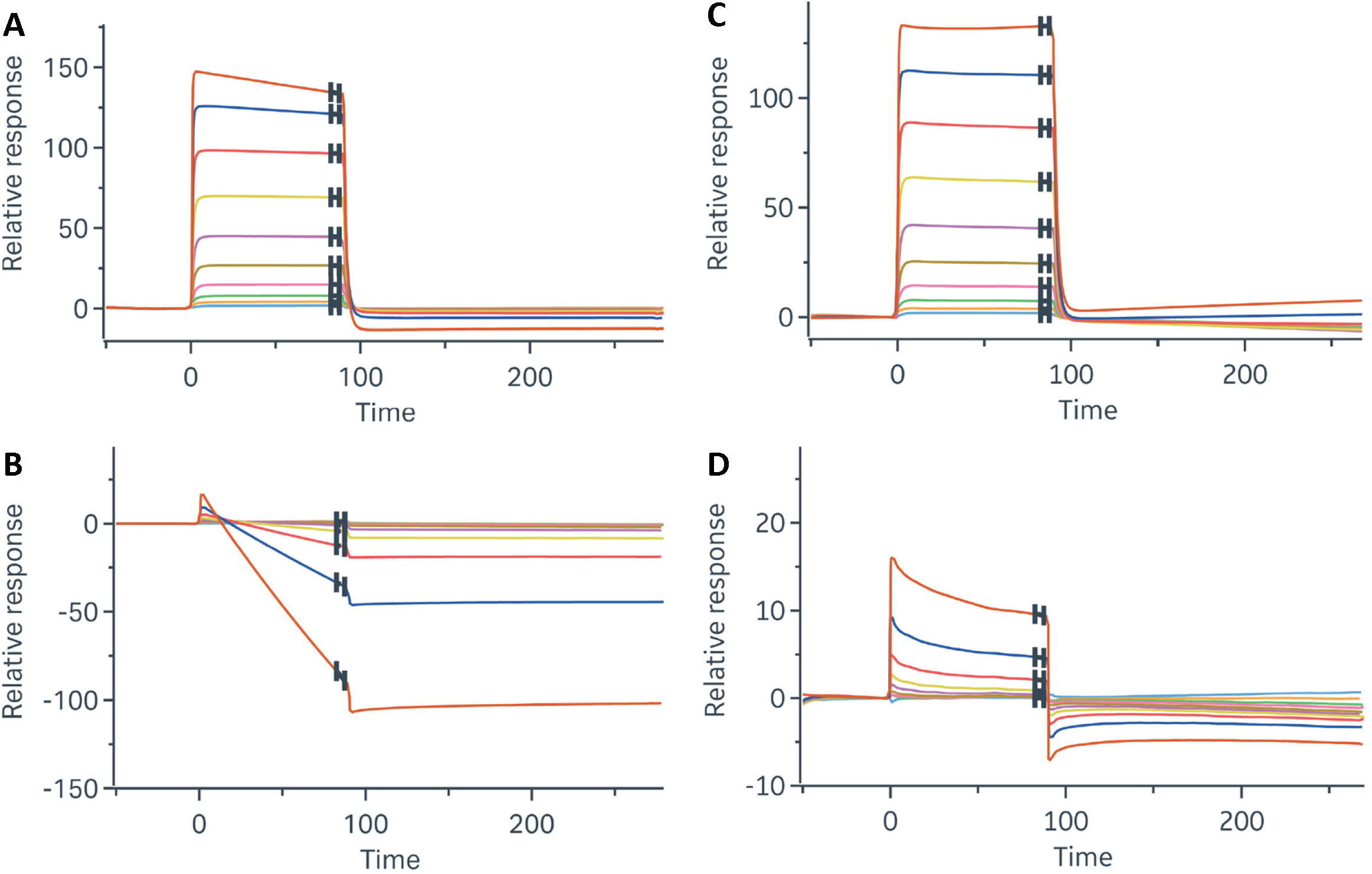
PYR exhibits modest glue activity for the DLG^W24C^-Kelch interaction as assessed by SPR analysis. SPR sensograms showed that the DLG^WT^-Kelch interaction displayed normal binding (A), whereas the DLG^W24C^-Kelch interaction was weakened due to the mutation (B). Upon PYR treatment, the DLG^WT^-Kelch interaction remained largely unchanged (C), while the DLG^W24C^-Kelch interaction showed a modest increase in binding (D).

### PYR binds KEAP1 possibly through a pocket in Kelch

To investigate how PYR promoted the interaction between NRF2^W24C^ and KEAP1, we performed ITC analysis and found that PYR bound to recombinant human KEAP1 with moderate affinity (Ka=7.54 µM, Kd = 13.27 µM, ΔH = -28.73kJ/mol, ΔS = -2.99 J/mol·K). These results suggested that PYR’s glue activity may depend on its direct binding to KEAP1. We then employed DiffDock to explore potential PYR-binding sites on KEAP1. Four candidate sites were identified: the BTB domain, a pocket within the Kelch domain, a surface pocket, and the DLG-binding site (Figure S5). Among these, the pocket within the Kelch domain exhibited the highest confidence score and the most consistent pose predictions across multiple runs (Figure S5B). This pocket also displayed a geometry favorable for forming a ternary complex, supporting its potential role in PYR’s glue activity. Molecular dynamics simulations further revealed that, compared to the DLG^WT^-Kelch interaction (Figure 5A), the DLG^W24C^-Kelch interaction showed a significant reduction in both interaction surface area and shape score, accompanied by an increase in binding energy (Figure 5B), indicating weaker and looser binding. However, upon docking PYR into the Kelch pocket, both the interaction surface area and shape score markedly increased - surpassing those observed in the DLG^WT^-Kelch complex - and the binding energy significantly decreased (Figure 5C). A 2D interaction map revealed that Arg^415^, Ala^556^, Gly^364^, Ile^416^, and Val^418^ interact directly with PYR (Figure 5D). These results suggested that PYR enhanced the interaction between NRF2^W24C^ and the Kelch domain by binding within this pocket.

**Figure 5.**
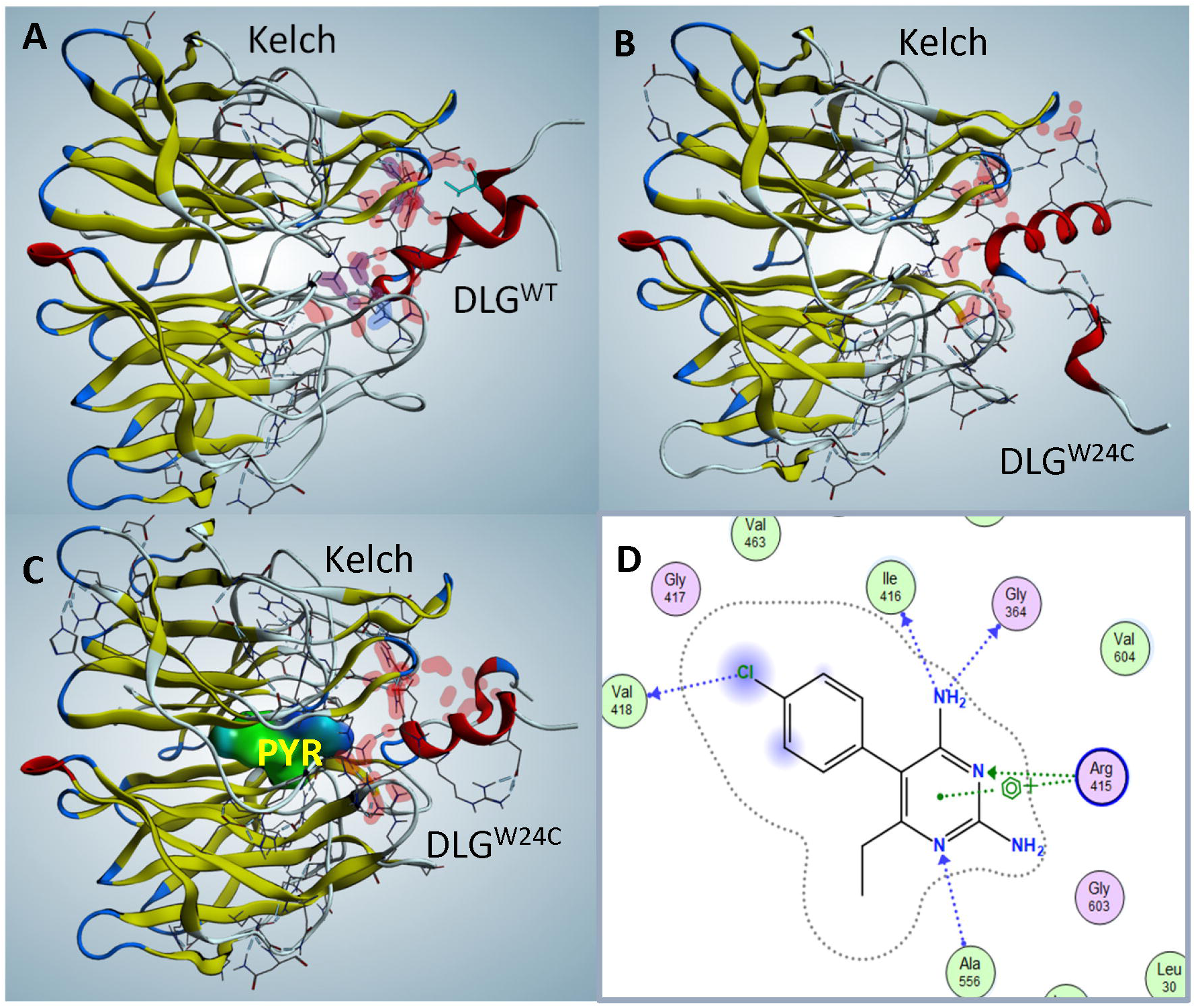
Molecular modeling suggested that PYR enhanced the NRF2^W24C^-KEAP1 interaction by binding a pocket within the Kelch domain. (A) The DLG^WT^-Kelch interaction involves key contacts between DLG motif residues (Asp^29^, Asp^27^, Arg^25^) and Kelch domain residues (Arg^380^, Arg^415^, Arg^483^). (B) The DLG^W24C^- Kelch interaction is weakened, indicating a looser binding interface compared to NRF2^WT^. (C) Incorporation of PYR into the Kelch pocket improved the DLG^W24C^-Kelch interaction, restoring surface complementarity. (D) A 2D ligand interaction map illustrated PYR’s contacts with residues in the Kelch pocket, including Arg^415^ and surrounding amino acids.

Interestingly, several known compounds have been reported to interact with Arg^415^ while disrupting the NRF2-KEAP1 interaction, thereby functioning as NRF2 activators [17–19]. Molecular docking of these compounds demonstrated that their carboxyl groups form strong ionic interactions with Arg^415^, occupying the canonical binding interface used by DLG or ETGE motifs to engage KEAP1. These compounds are relatively large in size and sit at the interface between KEAP1 and NRF2, effectively blocking their interaction (Figure S6). In contrast, PYR binds to a distinct site - within the Kelch pocket and opposite to Arg^415^ - and appears to pull Arg^415^ inward, thereby facilitating rather than disrupting the NRF2-KEAP1 interaction.

## Discussion

This study provides new insights into the functional consequences of NRF2 activation in ESCC and establishes PYR as a dual-mode NRF2 inhibitor. Using isogenic ESCC cell lines with genetically engineered deficiencies in NRF2 or KEAP1, we demonstrated that NRF2 activation produced broadly similar, yet context-dependent, effects on cellular behaviors, chemosensitivity, and radiosensitivity, varying according to the specific genetic context. A central finding of this work is the identification of PYR as a modest molecular glue that promotes the interaction between NRF2^W24C^ and KEAP1, but not NRF2^D29H^, NRF2^G81S^, or NRF2^WT^ (Figure 3, S4). This glue activity appears to depend on KEAP1. Molecular docking and dynamics simulation revealed that PYR binds a unique pocket within the Kelch domain, stabilizing the NRF2^W24C^-KEAP1 interaction. Thus, targeting KEAP1 with PYR led to NRF2 degradation.

Other reported NRF2 degraders, VVD-130037 and VVD-065, act differently by binding the BTB domain of KEAP1 and promoting KEAP1-CUL3 interactions [11]. Unlike these, PYR targets the Kelch domain offering a mechanistically distinct strategy for NRF2 degradation. Interestingly, VVD-065 altered the BTB homodimer structure in the opposite direction to NRF2 activators such as CDDO compounds, despite both targeting the same Cys^151^ residue. This bidirectional modulation of the BTB domain changes KEAP1-CUL3 affinity, enabling KEAP1 to sense both inhibitory and activating chemical signals (personal communication with Dr. Masayuki Yamamoto). In our study, molecular docking showed that PYR (Figure 5) and three NRF2 activators (Figure S6) interact with the same Arg^415^ residue in the Kelch domain. Analogous to the BTB domain, this bidirectional regulation of the Kelch domain fine-tunes NRF2-KEAP1 affinity, producing opposite effects on NRF2 activity. Future structural validation by X-ray crystallography or cryoEM will be essential to define PYR’s binding mode and the conformational dynamics of the KEAP1-PYR-NRF2^Mut^ ternary complex.

It is important to note that DHFR remains the primary target for PYR’s NRF2-inhibitory activity [14]. Through reduced folate downstream of DHFR inhibition, PYR can also inhibit STAT3 [21]. Consistently, PYR inhibited STAT3 signature genes in chronic lymphocytic leukemia patients with a good therapeutic response in a clinical trial [22]. However, systemic toxicity from DHFR inhibition [23] and its activity against both NRF2^Mut^ and NRF2^WT^ (Figure 3E, F) limit the clinical window for PYR. Identification of a unique pocket in the Kelch domain as a viable binding site for PYR underscored the potential of optimizing PYR’s glue activity and designing more potent glue, yet without the DHFR-inhibitory activity. Beyond NRF2 and STAT3, PYR has been shown to reduce the protein level of AIMP2-DX2, without affecting mRNA [15]. Previous studies also reported that PYR inhibits oncogenic proteins such as NFκB, AIMP2-DX2, MAPK, thymidine phosphorylase, telomerase, and others in various cancer cells [24]. Whether these effects result from direct binding or indirect DHFR inhibition remains unclear.

Another interesting observation in this study is that NRF2^WT^ and NRF2^Mut^ exhibiting distinct effects on cellular behaviors of ESCC cells. NRF2^WT^ overexpression in KEAP1^null^-KYSE450 cells promoted squamous differentiation. In contrast, NRF2^W24C^ expression exerted mixed effects on the expression of squamous differentiation markers. NRF2^W24C^ expression enhanced cell proliferation in KYSE70 cells, whereas NRF2^WT^ overexpression inhibited proliferation in KYSE450 cells (Figure 1). This observation is consistent with an *in vivo* mouse study in which concomitant expression of NRF2^L30F^ and TRP53^R172H^ resulted ESCC-like lesions. In contrast, while KEAP1^null^ induced similar NRF2 activation, concomitant expression of KEAP1^null^ and TRP53^R172H^ failed to generate ESCC-like lesions, and KEAP1^null^ cells disappear from the esophageal epithelium over time. These findings demonstrate that NRF2^Mut^ and NRF2^WT^ expression can elicit distinct fates of esophageal squamous epithelial cells [25]. Similarly, overexpression of NRF2^WT^ in KYSE410 and KYSE450 ESCC cells suppressed proliferation of ESCC. While ablation of NRF2^Mut^ in KYSE70 and KYSE180 cells caused decreased cell proliferation, expression of NRF2^Mut^ (NRF2^R34Q^ and NRF2^E79K^) accelerated the cancer cell proliferation [26]. These data indicate that NRF2^WT^ may act as a tumor suppressor in ESCC, whereas NRF2^Mut^ serves as an oncogene, underscoring the need for NRF2^Mut^-specific inhibitors over pan-NRF2 inhibitors like PYR.

In conclusion, this study broadens our understanding of PYR’s dual role in inhibiting NRF2. PYR acts both as a DHFR inhibitor and as a modest KEAP1-targeting molecular glue. The identification of a non-canonical glue site in the Kelch domain opens new opportunities for developing next-generation molecular glues to selectively target NRF2^Mut^-driven cancers.

## Acknowledgment

This work was supported by NIH R01 CA244236 and R01 AA030026 (to XC).

## Author’s contributions

Conceptualization: XC

Methodology: ZX, CP, HW, BS, CBL, YL, HL

Funding acquisition: XC

Writing original draft: ZX, XC

Writing, review & editing: ZX, XC

---

**Supplementary Table S1.** Antibody information.

| Antigen | WB | PLA | Co-IP | Catalog No. | Host | mAb/pAb | Company | City, State |
| --- | --- | --- | --- | --- | --- | --- | --- | --- |
| NRF2 | 1:1,000 |  |  | 20733 | Rabbit | mAb | Cell Signaling | Danvers, MA |
|  |  | 1:200 |  | 16396-1-AP | Rabbit | pAb | Proteintech | Rosemont, IL |
|  |  |  | - | GTX103322 | Rabbit | pAb | GeneTex | Irvine, CA |
| KEAP1 | 1:2,000 |  |  | ab119403 | Mouse | mAb | Abcam | Waltham, MA |
|  |  | 1:200 | - | ab119403 | Mouse | mAb | Abcam | Waltham, MA |
| Cytokeratin 1 (CK1) | 1:2,000 |  |  | LS-B7799-50 | Rabbit | pAb | Lifespan<br>Biosciences | Seattle, WA |
| Cytokeratin 4 (CK4) | 1:2,000 |  |  | LS-C165630-400 | Rabbit | pAb | Lifespan<br>Biosciences | Seattle, WA |
| Cytokeratin 14 (CK14) | 1:2,000 |  |  | ab7800 | Mouse | mAb | Abcam | Waltham, MA |
| CAV1 | 1:3,000 |  |  | A19006 | Rabbit | mAb | ABclonal | Woburn, MA |
| COL17A1 | 1:3,000 |  |  | A4808 | Rabbit | mAb | ABclonal | Woburn, MA |
| SOX2 | 1:2,000 |  |  | ab97959 | Rabbit | pAb | Abcam | Waltham, MA |
| P63 | 1:2,000 |  |  | ab124762 | Rabbit | mAb | Abcam | Waltham, MA |
| PAX9 | 1:2,000 |  |  | 12847 | Rabbit | mAb | Cell Signaling | Danvers, MA |
| HES1 | 1:3,000 |  |  | 11988 | Rabbit | mAb | Cell Signaling | Danvers, MA |
| GAPDH | 1:20,000 |  |  | ab8245 | Mouse | mAb | Abcam | Waltham, MA |

## Supplementary Figures

**Figure S1.**
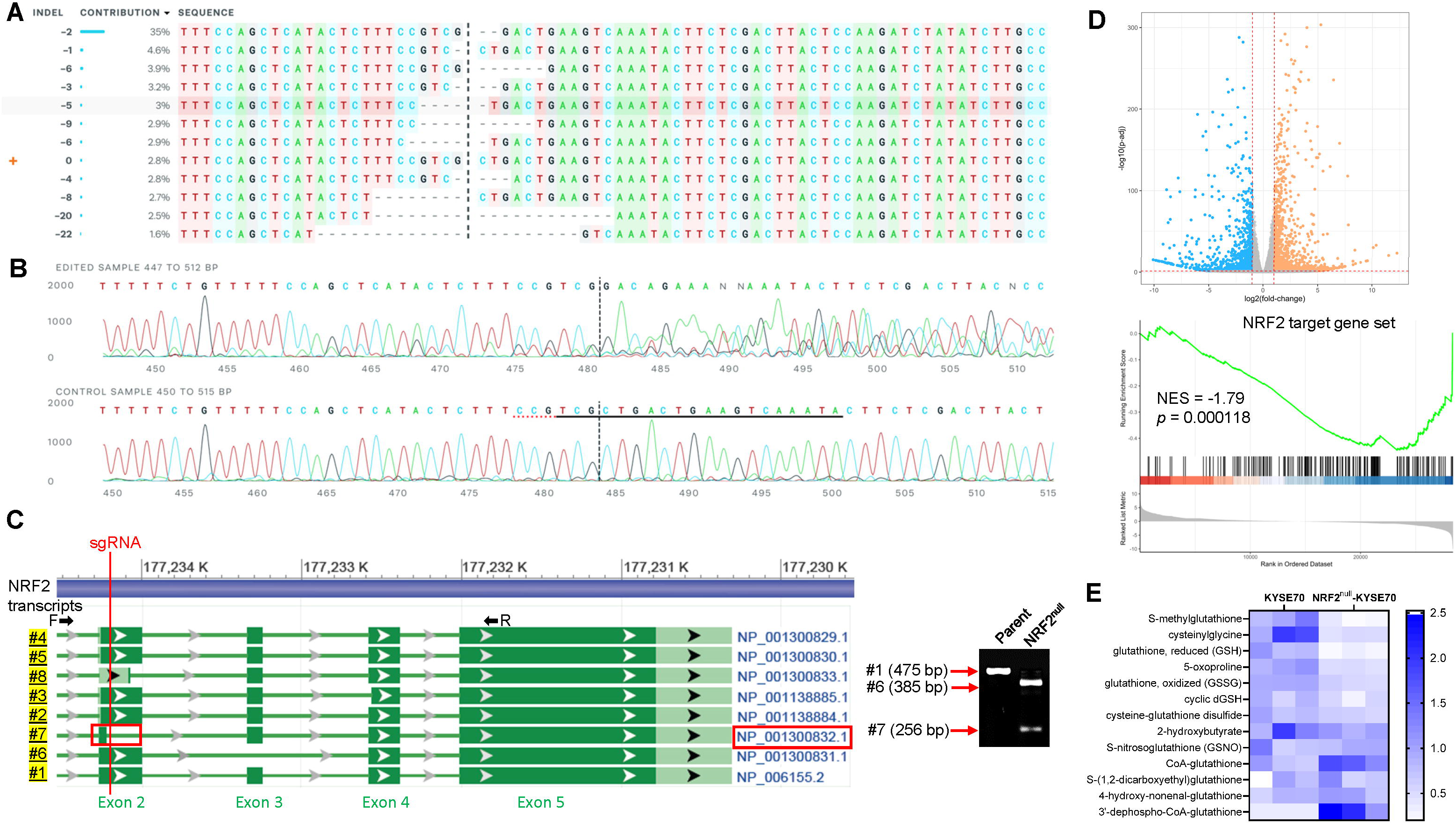
Isogenic NRF2^null^-KYSE70 cells were generated via CRISPR-Cas9 knockout for comparison with its parental NRF2^W24C^-KYSE70 cells. (A) Among the CRISPRed clones, -2 deletion was the most abundant one (35%) and chosen by monoclonal selection for subsequent experiments. The sgRNA target sequence for NRF2 was 5’-TATTTGACTTCAGTCAGCGA-3’. (B) Sanger sequencing validated 2-base deletion, which is expected to cause a frameshift mutation and a premature stop codon at amino acid 46 in exon 2. (C) A scheme of NRF2 transcript variants (#1-8) was generated according to NCBI. Vertical red line indicated that transcript 7 was missed by sgRNA. RT-PCR confirmed that transcript 6 (385bp) and transcript 7 (256bp) were expressed in the NRF2^null^-KYSE70 cells. Locations of forward and reverse primers were labeled. (D) Volcano plot of RNAseq data analysis and GSEA of NRF2^null^-KYSE70 cells in comparison to its parental NRF2^W24C^-KYSE70 cells showed that the human NRF2 target gene set was significantly downregulated in NRF2^null^-KYSE70 *vs* NRF2^W24C^-KYSE70 cells. (E) Metabolomic analysis showed that glutathione metabolism was significantly suppressed in NRF2^null^-KYSE70 *vs* NRF2^W24C^-KYSE70 cells.

**Figure S2.**
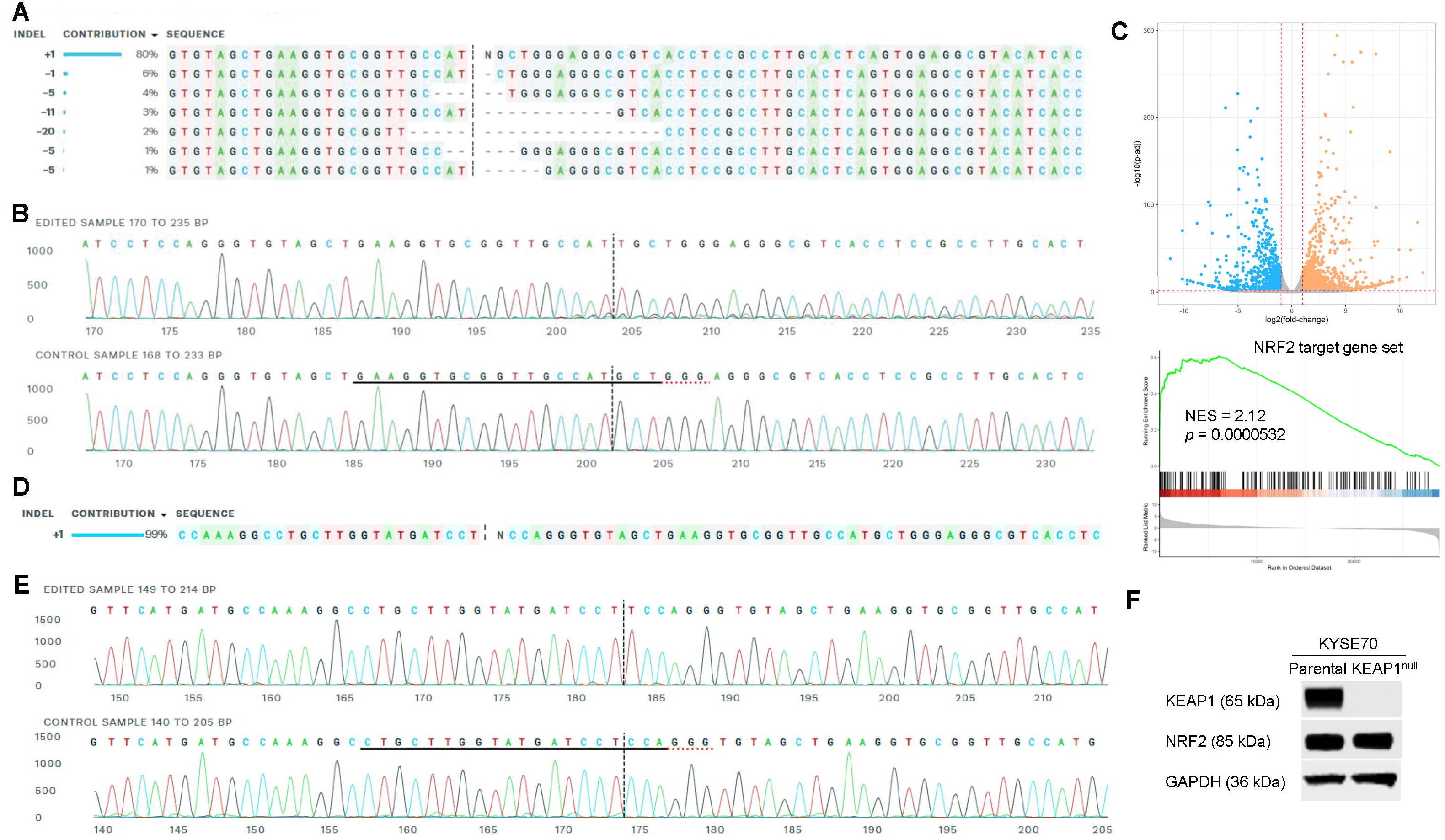
Isogenic KEAP1^null^-KYSE450 cells and KEAP1^null^-KYSE70 cells were generated via CRISPR- Cas9 knockout for comparison with their parental cells. (A) Among the CRISPRed clones of KYSE450 cells, +1 insertion was the most abundant one (80%) and chosen by monoclonal selection for subsequent experiments. The sgRNA target sequence for KEAP1 was 5’-GAAGGTGCGGTTGCCATGCT-3’. (B) Sanger sequencing validated +1 insertion in exon 2 which was expected to result in homozygous KEAP1 knockout. (C) Volcano plot of RNAseq data analysis and GSEA of KEAP1^null^-KYSE450 cells in comparison with its parental NRF2^WT^-KYSE450 cells showed that the human NRF2 target gene set was significantly upregulated in KEAP1^null^-KYSE450 cells *vs* NRF2^WT^-KYSE450 cells. (D) Among the CRISPRed clones of KYSE70 cells, +1 insertion was the most abundant one (99%) and chosen by monoclonal selection for subsequent experiments. The sgRNA target sequence for KEAP1 was 5’-CTGCTTGGTATGATCCTCCA-3’. (E) Sanger sequencing validated +1 insertion in exon 2 which is expected to result in homozygous KEAP1 knockout. (F) KEAP1^null^- KYSE70 cells were validated by absence of KEAP1 expression on Western blot.

**Figure S3.**
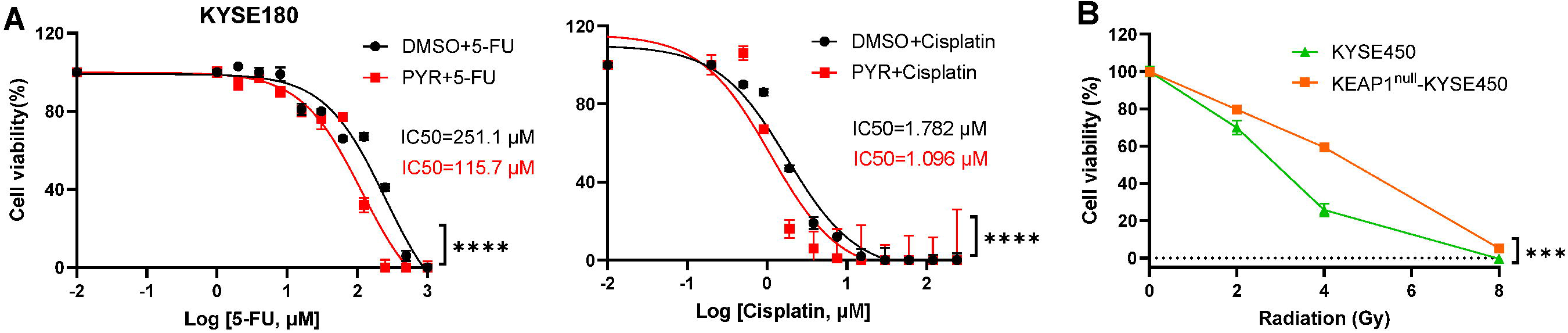
NRF2 modulates chemosensitivity and radiosensitivity of ESCC cells. (A) Viability of NRF2^D77V^-KYSE180 cells after co-treatment with PYR (10 μM) and 5-FU or cisplatin; (B) Viability of NRF2^WT^- KYSE450 and KEAP1^null^-KYSE450 cells in 3D culture when exposed to radiation. *** *p* < 0.001; **** *p* < 0.0001. Bar represented mean ± SD.

**Figure S4.**
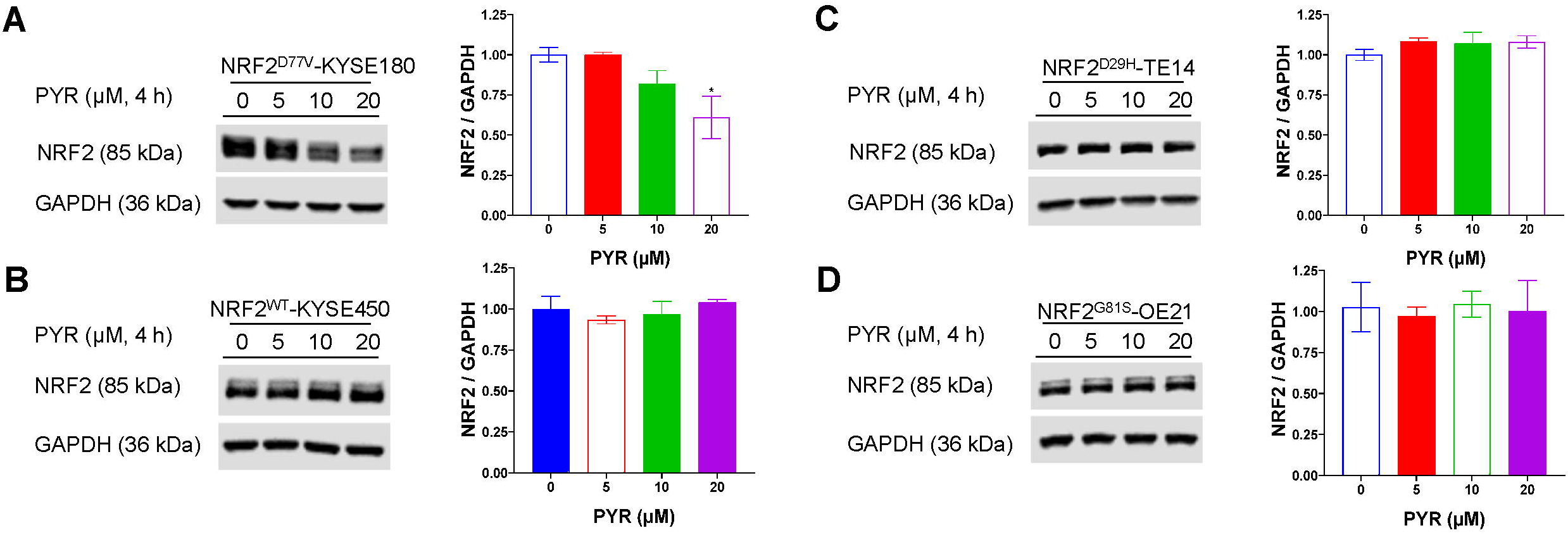
PYR had varied effects on NRF2 degradation. PYR (10 μM) degraded NRF2^D77V^ in KYSE180 (A) cells in a dose-dependent manner in 4 h, but not NRF2^WT^ in KYSE450 cells (B), NRF2^D29H^ in TE14 cells (C), or NRF2^G81S^ in OE21 cells (D). *\* p* < 0.05. Bar represented mean ± SD

**Figure S5.**
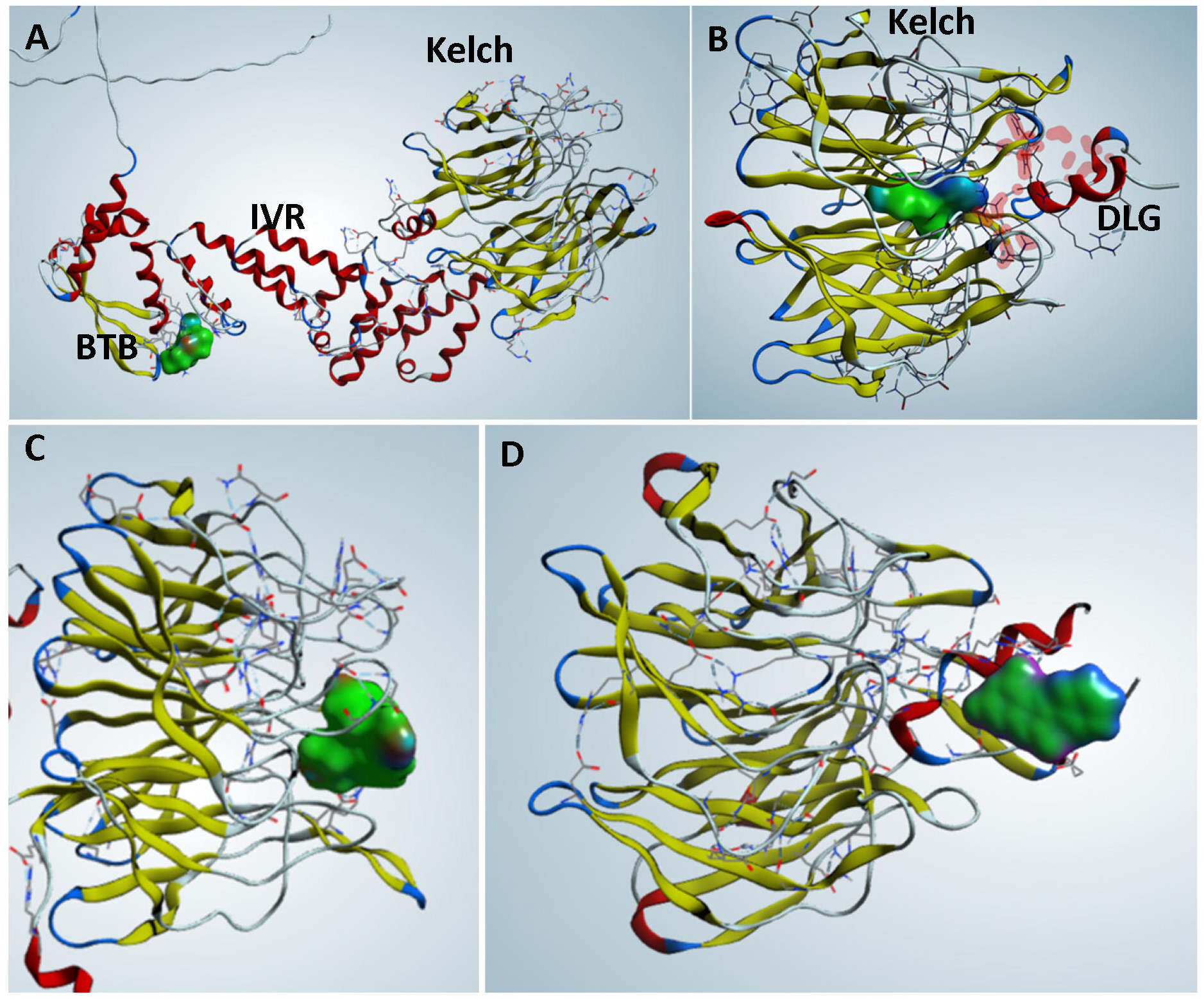
DiffDock identified four potential PYR-KEAP1 interaction modes.

**Figure S6.**
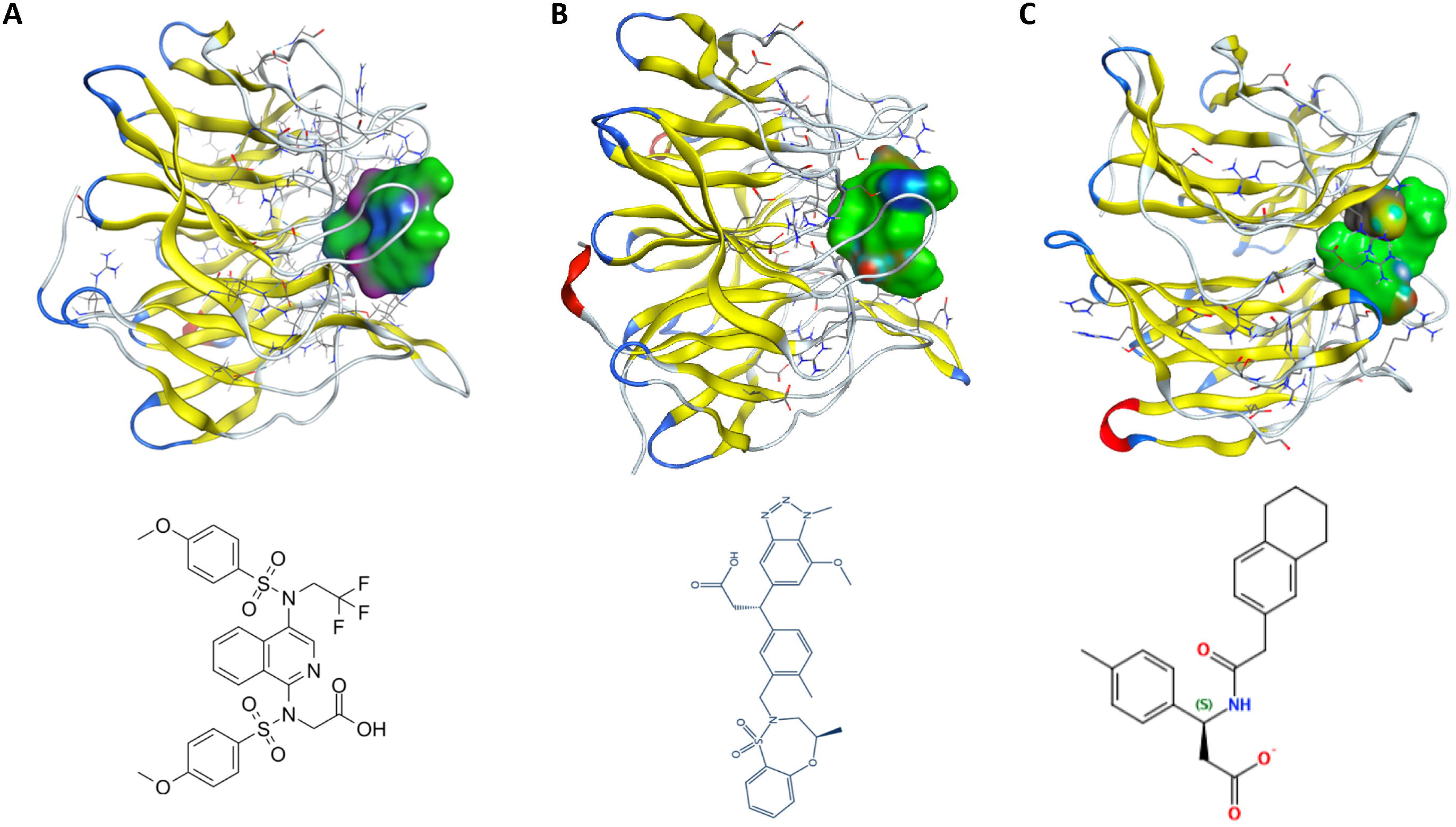
Compounds interact with Arg^415^ yet disrupt the NRF2-KEAP1 interaction and activate NRF2. (A) PRL-295-Kelch interaction; (B) KI696-Kelch interaction; (C) Compound 4-Kelch interaction.

## Supplementary Excel file

Excel S1. RNAseq data for differential gene expression in KYSE70 cells: NRF2^null^-KYSE70 *vs* parental NRF2^W24C^-KYSE70 cells.

Excel S2: RNAseq data for differential gene expression in KYSE450 cells: KEAP1^null^-KYSE450 *vs* parental NRF2^WT^-KYSE450 cells.

Excel S3. Metabolomics analysis of KYSE70 cells: NRF2^null^-KYSE70 *vs* parental NRF2^W24C^-KYSE70 cells.

## Notes

**Conflict of interest:** The authors declare no potential conflicts of interest.

### Competing Interest Statement

The authors have declared no competing interest.

